# Sensory Modality- and Load-Dependent Changes across Cortical Representations for Working Memory Storage

**DOI:** 10.1101/2024.02.02.578618

**Authors:** Vivien Chopurian, Simon Weber, Thomas B. Christophel

## Abstract

While distributed cortical areas represent working memory contents, their necessity for memory maintenance has been questioned. We examined the differential effects of maintaining multiple items on the neural information across cortical regions. In each trial of the fMRI experiment, participants (n=81) had to memorize two items, each either an orientation or a pure pitch, for 13.8s and continuously recalled the target after the delay. We kept the overall working memory load constant, but varied the sensory modality of each item to vary the effective visual load. We show that increasing visual load decreased behavioural recall performance for orientations and continuous orientation-specific decodable information in visual cortex but less so infrontoparietal areas. Simulations show that this drop in decodable information is best interpreted as a drop in mnemonic information represented by patterns of visual cortex activity. Our results provide evidence for shared labour of visual cortices, where maintaining two versus one orientation leads to a loss in representational fidelity, and anterior cortices, where multiple items could be represented in a more robust but less precise format.

## 1 Introduction

Working memory capacity is limited. The precision of recall decreases as we increase the number of items to remember (Gorgoraptis et al., 2011; Ma et al., 2014). In neuroimaging studies, the load dependent decrease in recall performance is accompanied by a decline in the strength of visual working memory representations (Emrich et al., 2013; Sprague et al., 2014; Sutterer et al., 2019). While this evidence demonstrates a link between the ability to decode cortical representations and the capacity to memorize information, it remains unclear whether different cortical regions play distinct roles as memory load increases.

Previous studies highlight a distributed network of sensory, parietal and frontal regions to maintain memorized contents during working memory delay (Christophel et al., 2017; D’Esposito, 2007; Fuster, 1998; Xu, 2023). For single visual items, studies have found correlations between decodable information in visual cortex and recall performance across participants and trials (Ester et al., 2013; Hallenbeck et al., 2021; Weber et al., 2024). The neuronal selectivity in visual cortex might enable the maintenance to sensory-like representations of memorized content (Ester et al., 2013; Harrison & Tong, 2009; Serences et al., 2009). These representations might be modulated by incoming visual input, as they have been found to be susceptible to visual distractors, leading to reduced delay-period decoding if the distractors also affected working memory recall (Hallenbeck et al., 2021; Lorenc et al., 2018; Rademaker et al., 2019). Imaging and single-cell studies demonstrated maintenance of item-specific representations in parietal and prefrontal regions (Bettencourt & Xu, 2016; Ester et al., 2015; Funahashi et al., 1989; Goldman-Rakic, 1995) and persistent delay activity in prefrontal cortex directly correlated with behavioural performance in non-human primates (Constantinidis et al., 2001; Wimmer et al., 2014). Presumably, anterior regions maintain abstracted, sensory-modality independent mnemonic representations (Leavitt et al., 2018; Spitzer & Blankenburg, 2012; Vergara et al., 2016), which appear to be protected from interference during visual distraction (Bettencourt & Xu, 2016; Lorenc et al., 2018; Rademaker et al., 2019). Furthermore, frontoparietal regions maintained information for multiple items independent of priority, whereas only the prioritized item was decodable from the visual cortex (Christophel et al., 2018). Taken together, visual cortex as well as frontoparietal regions have been found to maintain item-specific mnemonic information, but the difference in distractor-susceptibility across these regions could suggest different roles for working memory storage of multiple items.

If multiple low-level visual items have to be encoded and maintained in working memory, their representations are likely to compete for resources in visual cortex (Franconeri et al., 2013). Prior work demonstrates that the extent of cortical overlap of multiple representations predicts the degree to which memory recall is impaired (Cohen et al., 2014). Therefore, we assume that working memory contents from different sensory modalities should interfere less because these representations share little overlap within sensory regions (Baddeley, 2003; Kiyonaga & D’Esposito, 2020). The recruitment of abstract anterior representations (Christophel et al., 2017; Chunharas et al., 2023) could reduce interference by helping to separate similar sensory representations, but perhaps at the cost of precision (Bae, 2021).

In this study, we investigated the effect of visual working memory load on these distributed cortical representations. We show, by keeping the overall working memory load constant at two items but varying the sensory modality of each item, that increasing visual working memory load selectively reduced continuous mnemonic information about items present in visual cortex (V1-V3) but less in intraparietal sulcus (IPS) and precentral sulcus (sPCS).

## 2 Methods

### 2.1 Participants

Eighty-six participants were scanned. Four data sets with incomplete recordings, resulting from participant dropouts, and one data set with image artifacts were removed from the analysis. Our final sample consisted of 81 participants (48 female, 4 left handed, *mean*_*age*_ = 25, *sd*_*age*_ = 4). All participants provided their written informed consent and were compensated with 10€ per hour for their participation. The study was approved by the ethics committee of the Humboldt Universität zu Berlin (2022-40) and conducted according to the principles of the Declaration of Helsinki (World Medical Association, 2013).

### 2.2 Task and Procedure

In each trial of the delayed-estimation task, participants had to memorize two sequentially presented items (0.4s, inter-stimulus interval 1s), either two visual orientation stimuli, two auditory pitch stimuli or one of each. Crucially, this means that overall working memory load was constant at two items, but unisensory visual load varied between one or two items (see Figure 1). We focused all our analyses on decoding visual stimuli but included the auditory-auditory condition (auditory Load 2) to balance the design behaviourally. Whenever a sample of a given modality was presented, the other modality was masked. After an extended delay (13.8 s), a visual retro-cue (‘1’ or ‘2’ for 1.2s) indicated whether the first or second item had to be recalled. During the continuous recall phase (4.8s), participants had to adjust a probe orientation or pitch to match the target item by pressing the inner left and right button, and after confirming their response the fixation dot turned green. The next trial started after a brief period of fixation (1.6, 2.4 or 3.2s).

**Figure 1.**
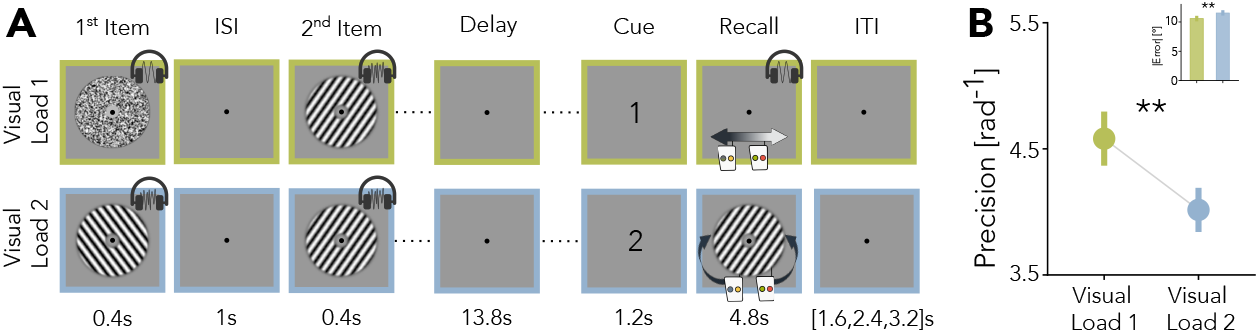
Task setup and behavioural results. **A** Example experimental trials with varying visual load. The top row shows a visual Load 1 trial, where items from both modalities are shown and the target modality is auditory. The bottom row shows a visual Load 2 trial, where both items are visual. Auditory Load 2 trials (not shown) are included in the design to balance the behavioural task. Visual samples were presented jointly with auditory noise and auditory samples were presented with visual masks. **B** Behavioural results for visual targets. Inset shows absolute error for the visual load conditions. Asterisks indicate significance levels of \*\**p <* .01. Error bars show the SEM.

The experiment consisted of 192 trials, divided in four runs with short breaks in between. The item modality was counterbalanced across cued and non-cued items and 12 orientations and pitches were counterbalanced for each modality and randomized across each cued and non-cued items (see Stimuli). Thus, the two orientations in each load condition were independent from one another. We randomized the experimental conditions within each scanner run to minimize effects of biased training sets for the cross-validation procedure (see fMRI Analyses). Before the main experiment, participants read instructions presented inside the scanner and completed 12 training trials during an anatomical scan. During these training trials, participants received feedback about their performance as the difference between their response and the target item.

### 2.3 Stimuli

Stimuli were generated with MATLAB and Psychtoolbox-3 (Brainard & Vision, 1997). The background color was set to grey (RGB [128,128,128]). The fixation dot (radius = 0.1°) was centered on the screen. Orientation stimuli and masks were presented as annuli with a total diameter of 7°, an inner diameter of 1.5° and a gaussian blur (gaussian kernel = 1°, sd = 0.5°) on all edges. The visual stimuli consisted of 12 discrete orientations with a 15° distance, from 7.5° to 172.5° with a 15° distance, excludeding the cardinal axes. The spatial frequency was set to one cycle per degree. Visual masks were generated by filtering white noise to only include spatial frequencies between 1 and 4 cycles per degree (code from (Rademaker et al., 2019)). During the recall, participants could select their response out of orientations between 1° to 180°, with a precision of 1°. For the auditory stimuli we used 12 pure tones with a distance of one semitone, between 251.63Hz and 493.88Hz, with a logarithmic ramp up and down of 0.02 seconds. White noise was chosen as auditory masks. The volume was adjusted for participants comfort level. During the recall period, a random pitch was played, and participants could select a response between 246.93Hz and 523.24Hz, with a precision of *Frequency ×* 2^±1/(70/1)^. During the recall phase, a longer button press increased the response step size, making it easier for participants to select their response.

### 2.4 Magnetic resonance imaging

Imaging data was recorded on two 3-Tesla Siemens Prisma MRI scanners with a 64 channel head coil. The first 40 participants were scanned at the Berlin Center for Advanced Neuroscience (Charité Berlin, Germany) with the Syngo MR VE11C operating system, a viewing distance of 148cm and screen of dimensions of 52 × 39cm. The subsequent 41 participants at the Center for Cognitive Neuroscience Berlin (Freie Universität Berlin, Germany) with the Syngo MR XA30 operating system, a viewing distance of 106cm and screen of dimensions of 48 × 27.2cm. Stimulus size was adjusted according to the screen-size and viewing distance, so that all participants received the same visual input. The decoding results for both scanners did not differ qualitatively from the combined results. At the beginning of each session, we recorded T1-weighted MPRAGE structural image (208 sagittal slices, TR = 2400 ms, TE = 2.22ms, flip angle = 8°, voxel size = 0.8mm^3^ isotropic, field of view = 240*256 mm). For each of the four runs, we recorded functional images with a multi-band accelerated echo planar imaging sequence, with a multi-band factor of 8 (TR = 800ms, TE = 37ms, flip angle = 52°, voxel size = 2 mm^3^ isotropic, 72 slices, 0mm inter-slice gap, field of view = 1872mm).

### 2.5 Preprocessing

All preprocessing steps and analyses were performed in MATLAB, using SPM12 (Ashburner et al., 2014), The Decoding Toolbox Version 3.999E2 (Hebart et al., 2015), and in-house software. First, we converted the acquired DICOM images into NIfTI format. The intensity-normalized anatomical image (T1) was coregistered to the first functional image of the first run for each participant. During segmentation, we estimated the normalization parameters to the Montreal Neurological Institute (MNI) standard space. Functional images were spatially realigned to the first image of the first run and resliced. To remove slow signal drifts accumulating across each run, we applied a cubic spline interpolation to each voxel’s time series (n_nodes_ = 24). Afterwards, we applied temporal smoothing by running a moving average of 5 TRs to the data of each run. We used the detrending algorithm and moving average as described in (Weber et al., 2024).

### 2.6 Identifying ROIs

We focused on the distribution of representations across three regions of interests (ROI) using probabilistic maps of retinotopic regions of the human neocortex (Wang et al., 2015). To generate a mask for visual cortex, we combined maps for areas V1 to V3 in standard space. To obtain the bilateral IPS mask, we combined IPS1 to IPS6 from the same probabilistic atlas, and similarly, for the sPCS left and right hemispheres were combined into one mask. After transforming these maps into single-participant space by applying the inverse normalization parameters estimated during preprocessing, we excluded voxels that had less than a 10% probability to belong to each ROI respectively. Secondly, we obtained activity driven maps by estimating a participant-level univariate GLMs for onsets of every orientation presented. After contrasting visual (orientation) activation and baseline, we selected the 1000 most active voxels within the participant-level anatomical ROIs. We selected univariate activated voxels, as previous work suggests that multivariate item-specific effects may emerge because of voxel-wise item-selective univariate signals (Albers et al., 2018; Haynes, 2015).

### 2.7 fMRI Analyses: pSVR

We analyzed fMRI data using support vector regression (SVR) (Chang & Lin, 2011) to continuously reconstruct memorized orientations. The orientation labels were transformed from the 0°-180° space into radians from −pi to pi. The angular orientation labels (*θ*) were then projected into sine and cosine labels. This provided a linear measure of these orientations and we predicted each set of these new labels from the multivariate voxel pattern of each ROI with a SVR with a radial basis function kernel (periodic SVR, (Weber et al., 2024)). In each cross-validation fold, we rescaled the training data voxel activation to 0-1 and applied the estimated scaling parameters to the test data. The orientation label was reconstructed by using the four-quadrant inverse tangent:

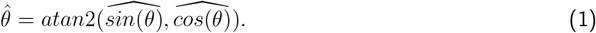

We calculated the circular absolute difference between presented and predicted orientation label:

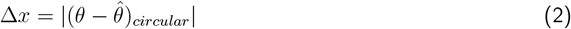

and this difference was then transformed into a trial-wise feature continuous accuracy (FCA) above chance to make our results comparable to existing literature applying this measure (Degutis et al., 2024; Weber et al., 2024) as well as to classification-based measures.

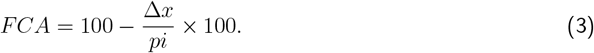

We subtracted 50% from the resulting FCA, so that 0% refers to chance level decoding with an error of 45°. This conversion means that values larger than chance level imply a smaller decoding error.

All pSVR analyses were run for each TR individually and cross-validated across the four runs. On each cross-validation iteration, we trained on the trials of the three runs and tested on the left-out run. In order to increase computational power, we combined all orientations within each of the load conditions (Figure 2A). Thus, we ran separate decoding schemes for condition Load 1 and Load 2. For the temporal generalization analysis (Figure 4), we trained the pSVR on each time-point and tested on each timepoint. This was done for each Load condition separately as well as across Load conditions, where we, for instance, trained on each time point from Load 1 and tested on each time point from Load 2.

**Figure 2.**
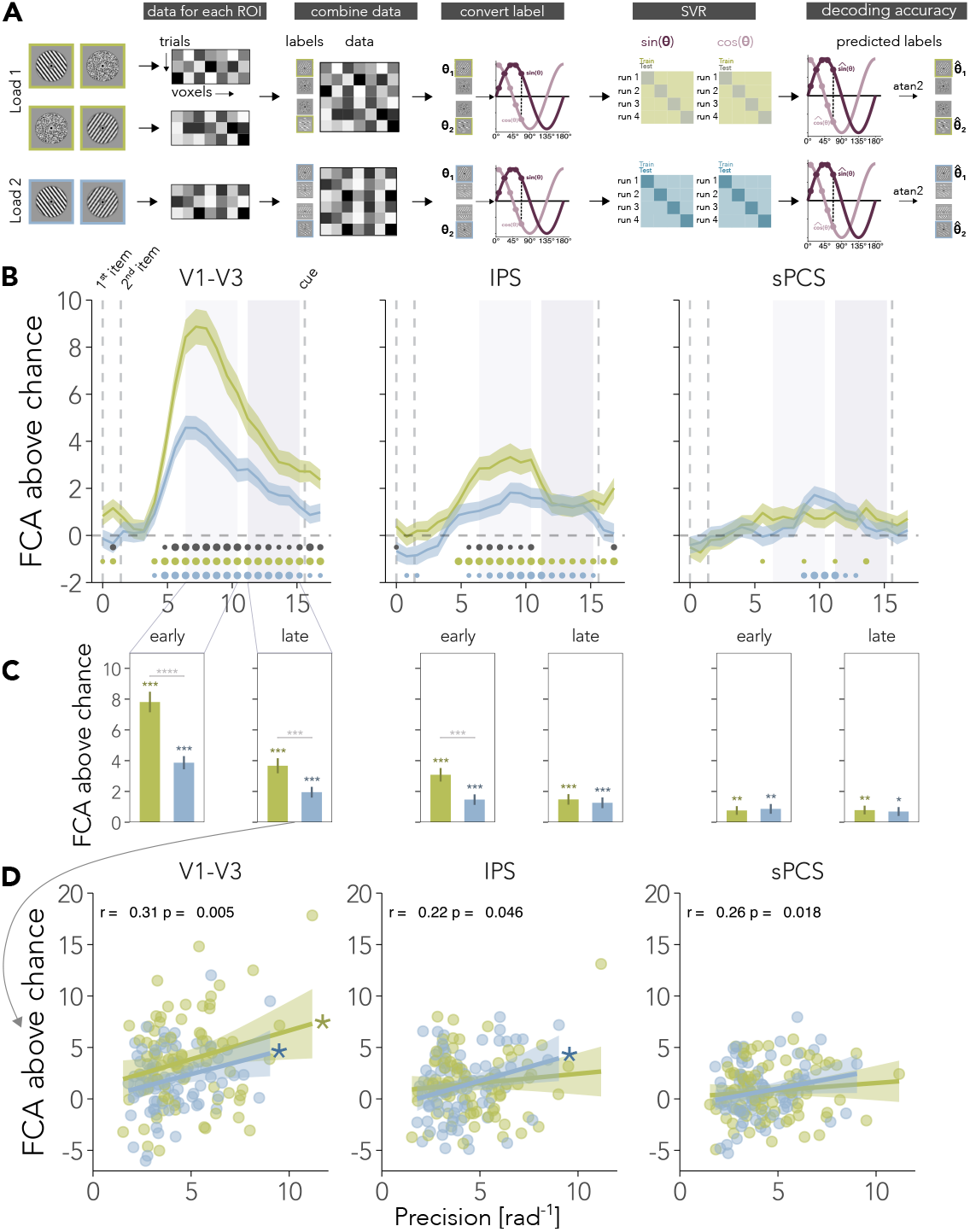
Decoding memorized orientations across cortical regions. **A** Analysis steps for each ROI and TR. We converted the orientation labels into sine and cosine labels. A support vector regression (SVR) was applied to predict the sine and cosine label separately (Weber et al., 2024). The difference between the orientation labels and predicted labels was converted into feature continuous accuracy (FCA). An FCA of 0% refers to chance level decoding error of 45°. See Methods for details. **B** Time-course of item-specific decoding accuracy for each visual Load condition in all three ROIs. Early (6.4s-10.4s), where signals can be attributed to perceptual and mnemonic signals, and late delay (11.2s-15.2s), where signal can be attributed to working memory maintenance, were chosen to avoid overlap.**C** Average difference between Load 1 and Load 2 in early and late delay. Asterisk above each bar indicate uncorrected above chance significance. **D** Correlations between late delay FCA and working memory recall precision. Correlation coefficient in figures refers to correlation across both conditions. Dots (small, medium, large) at the bottom of **B** and asterisks in **C** indicate significance levels of \**p <* .05, \**p <* .01 or \*\*\**p <* .001. In **B**,**C** shaded ribbons and error bars show SEM and 95% confidence interval in **D**.

For Figure 3C, we simulated a load 1 conditions across different signal strength levels. To evaluate how training and testing on both orientation labels in load 2 impacts the decoding accuracy, we compared our analysis strategy of decoding both items jointly to a condition, where the first and second item are decoded separately.

**Figure 3.**
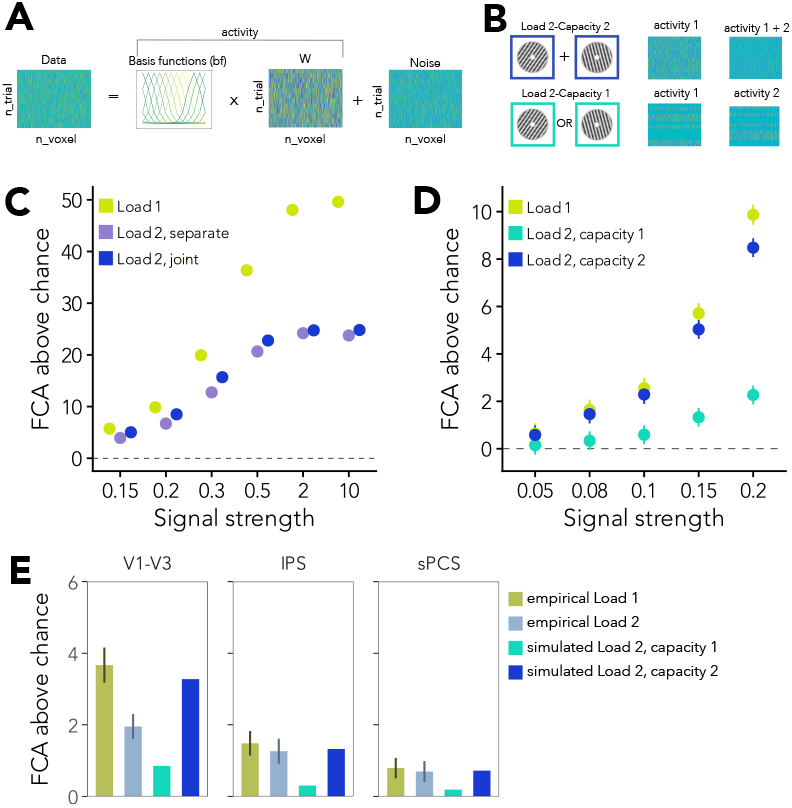
Simulated data show differences between decoding schemes and capacity limits **A** Schematic simulation pipeline. **B** Multivariate pattern for Load 2 conditions. Addition of activity patterns for Load 2-Capacity 2 or random selection of a single pattern per trial for Load 2-Capacity 1. **C** Results for our simulated Load 1 condition and two Load 2 condition with different decoding schemes. Load 2 - separate refers to the separate decoding of the two orientation labels, while Load 2 joint refers to the decoding scheme used in our experiment, where we train and test on both labels. **D** FCA above chance for simulated load conditions across different signal strength levels. **E** Empirical results for Load 1 and 2 in late delay period and empirical Load 1 divided by ratios for simulated load conditions for each ROI. Errorbars show SEM.

**Figure 4.**
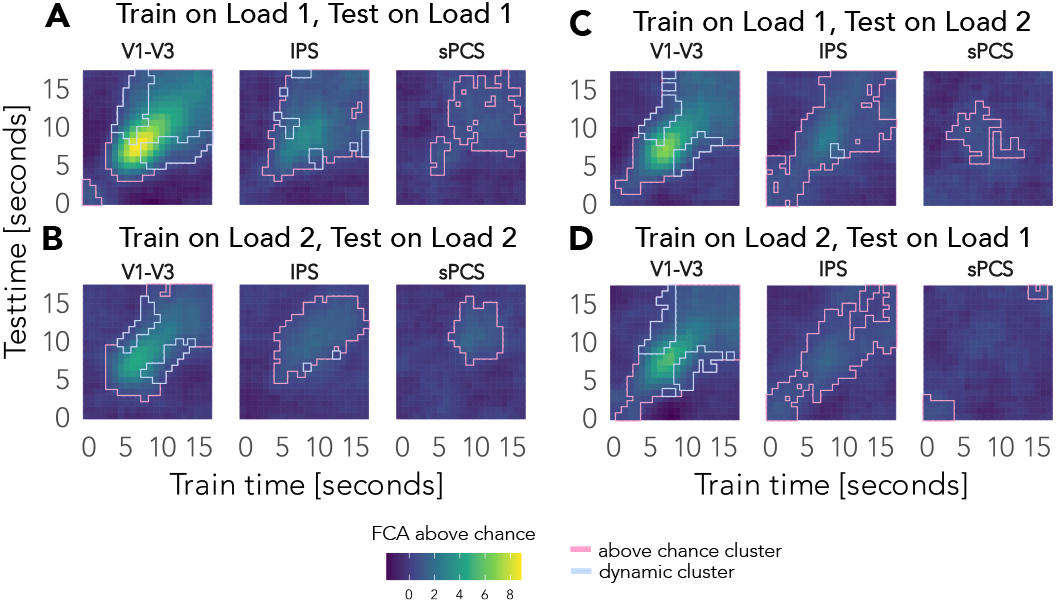
Mnemonic representations generalize between conditions in visual and parietal areas Results for cross-condition temporal generalization, where we trained on each time-point in each load condition and tested on each time-point in the other load condition. Cross-temporal decoding for **A** Visual Load 1 and **B** Visual Load 2. Cross-temporal decoding when **C** training on TRs from Load 1 and stesting on TRs from Load 2, and **D** training on TRs from Load 2 and testing on TRs from Load 1. White dotted lines indicate first item, second item and cue onset. Significant above chance clusters and dynamic clusters were tested for p *<* 0.05.

### 2.8 Statistical analyses

We analysed the behavioural data for the orientation recall by calculating precision (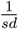 in radians) for the angular difference between target orientation and response orientation for each condition and participant(Gorgoraptis et al., 2011). Since the orientation space is circular, we used the circular SD. We compared the average of each visual load condition with paired two-sided t-tests.

To test for significant differences in decoding accuracy between the load conditions (visual Load 1 and 2), we used paired two-sided t-tests. In particular, we tested decoding accuracy during a period early in the delay, where we assume to find perceptual activation mixed with memory representations (due to hemodynamic lag), and a period later in the delay, where we assume to find information about mnemonic representations without a substantial influence of perceptual representations. We averaged decoding accuracy per participant in each of these two sections of the delay period (early delay: 6.4-10.4s; late delay: 11.2s-15.2s). Then we tested for the difference between the load conditions within each delay period with paired two-sided t-tests and within each condition for decoding accuracy above chance (50%) with two-sided t-tests. We focused all our other analyses on the late delay. Where applicable, for example when comparing results across ROIs and load conditions, we corrected for multiple comparisons with Bonferroni correction. For descriptive purposes, we also compared decoding accuracy to chance level and between conditions (Figure 2B) with two-sided t-tests for each time-point (22 TRs), from the onset of the first stimulus to recall onset. Both analyses were validated with leave one run out cross-validation across participants. Correlation coefficient refers to Pearson’s correlation. Correlations were compared with the R package cocor (Diedenhofen & Musch, 2015), where we report the p-values for Pearson and Filon’s z, and partial correlations were calculated with the R package ppcor (Kim, 2015)

### 2.9 Simulations

To determine how different analyses choices may influence the results, we simulated multivoxel data for load conditions 1 and 2 with varying levels of signal strength. We simulated data for a single time point with the same experimental parameters as in our fMRI experiment (n participants = 81, n runs = 4, n labels = 12, n voxels = 1000) and ran each simulation for 1000 iterations. First, we defined the similarity of the orientation space with 12 cosine functions raised to the power of 11 (Sprague et al., 2019), covering the whole orientation space (0°-180°) and their centers set at each stimulus label. Next, we defined a weight matrix (W) for each participant. This weight matrix models the voxel sensitivity for the orientation space and stayed constant for each participant throughout the iterations. The noise is white noise sampled from a normal distribution with mean *µ* = 0 and *σ* = 1 (Welvaert & Rosseel, 2014):

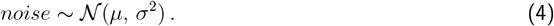

We simulated the different load conditions with different *noiseFactors* (e.g., 0.1, 5, 12.5) to a *signalFactor* of 1, which weights the noise and signal respectively, to approximate our empirical level of decoding accuracy. The simulated multivariate voxel pattern for a single orientation can thus be described as:

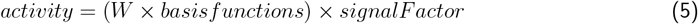

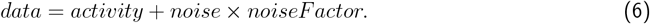

Next, to determine whether the drop in visual cortex decoding accuracy is evidence of visual cortex only maintaining one item in the focus of attention (Sutterer et al., 2019), we simulated two load 2 conditions.

First, we simulated the effect of additive data with increased working memory load (Load 2 - Capacity 2), where we generated the multivoxel pattern for each orientation label and then added two patterns, assuming a simplified superpositional code for sequentially presented items (Botvinick & Plaut, 2006):

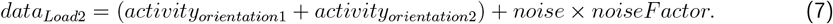

Additionally, we simulated a capacity of one by randomly selecting the pattern for one of the items in each trial before adding the two patterns (Load 2 - Capacity 1; see Sutterer et al. (2019)).

In our data, FCA drops from Load 1 to Load 2 by a factor of 2 in V1-V3. To investigate which simulated load 2 condition matches this ratio, we calculated the ratio of Load 1/Load 2 for each of our simulated load conditions and signal strength. For Figure 3D, we then divided our empirical FCA for Load 1 by the ratio based on signal strength for each delay period and ROI in Load 1.

**Table 1.** Simulated Load 1/Load 2 ratios compared to empirical data ^*a*^.

| Signal strength | sim. Load 2-Capacity 2 | sim. Load 2-Capacity 1 | Empirical data |
| --- | --- | --- | --- |
| 0.1 | 1.12 | 4.32 | V1-V3: 1.89 |
| 0.08 | 1.12 | 4.86 | IPS: 1.18 |
| 0.05 | 1.10 | 4.24 | sPCS: 1.14 |

<sup>a</sup> Signal strength refers to the ratio between different noise factors to a signal factor of 1.

### 2.10 Temporal generalization analysis

To assess whether the neural code is generalizable across time and load conditions, we conducted a temporal cross-decoding and cross-condition analysis. Here, we trained on data on one time point and tested on data from all other time points, within and across the load conditions (Figure 4). As in the main analysis, the pSVR was cross-validated by training on three runs and testing on the left-out run. We applied a cluster-based test (Maris & Oostenveld, 2007) to determine above-chance clusters, where the neural code was generalizable across time. First, we generated a null distribution by randomly assigning the sign (an FCA of 0 refers to chance level) of all elements of the decoding accuracy matrix. Then we calculated the summed t-value for the largest randomly occurring cluster and repeated this procedure 1000 times. The resulting null distribution cluster were then compared to our empirical summed t-value with a threshold of p *<* 0.05 to determine whether the size of the clusters were larger than chance (Degutis et al., 2024).

Dynamic clusters were elements of the temporal-cross decoding matrix where the neural code at one time point did not generalize to another time point, within-and across conditions. This was defined as off-diagonal elements which were significantly smaller than their two corresponding diagonal elements (*a*_*ij*_ *< a*_*ii*_ and *a*_*ij*_ *< a*_*jj*_). To test whether the accuracy was significantly smaller, each of these two conditions was tested with a cluster-based permutation test. We subtracted the diagonal element from the off-diagonal element (*a*_*ij*_ *− a*_*ii*_ and *a*_*ij*_ *− a*_*jj*_) and applied the same sign permutation test for both comparison as mentioned above. Only if both tests were significant (p *<* 0.05) and the element was part of the above-chance cluster, the element was defined as dynamic (Degutis et al., 2024; Li & Curtis, 2023; Spaak et al., 2017).

### 2.11 Bugs

For the first 11 participants, the number of images recorded in each run was set to 1448 instead of 1460, hence ending immediately after the last trial. Nineteen participants (pattern 1: 11, pattern 2: 8) had the same randomisation for the trial order due to a glitch in the code. The results did not change qualitatively after removing these participants from the analyses.

## 3 Results

### 3.1 Visual load reduces mnemonic information in visual cortex

On trials with visual recall, participants performed better on trials where only the target item was an orientation and the other item was a pitch (visual Load 1, *precision* = 4.70*rad*^*−*1^) compared to the visual Load 2 condition (visual Load 2, *precision* = 4.13*rad*^*−*1^, *t*_(80)_ = 3.06, *p* = 0.003, Figure 1B). To investigate this visual load effect on the distributed cortical representations, we decoded orientations continuously with a periodic support vector regression (pSVR Weber et al. (2024), see Methods and Figure 2A) from fMRI data. While all three regions of interest, the visual cortex (V1-V3), intraparietal sulcus (IPS) and precentral sulcus (sPCS), show information about orientations, the feature continuous accuracy (FCA) in each region changed with load and across time (Figure 2B).

The decoding accuracy in visual cortex was affected by visual load throughout the trial. Our results showed main effects of load (*F*_(1,80)_ = 47.57, *p <* 0.0001) and time (*F*_(1,80)_ = 54.46, *p <* 0.0001), as well as an interactions between load and time (*F*_(1,80)_ = 14.16, *p <* 0.001) on decoding accuracy. Early in the delay (6.4s-10.4s after first item onset) decoding accuracy decreased with increased visual working memory load (*t*_(80)_ = 7.53, *p <* 0.0001, Bonferroni corrected). In this early delay phase cortical representations can be assumed to be a mixture of perceptual and mnemonic representations due to the hemodynamic lag. However, the same pattern of results is also present late in the delay (11.2-15.2s after first item onset; see Rademaker et al. (2019)), when signals can be attributed only to working memory signals (*t*_(80)_ = 3.52, *p* = 0.002, Bonferroni corrected). While in both conditions decoding accuracy dropped from early to late delay (Load 1: *t*_(80)_ = 7.19, *p <* 0.0001, Load 2: *t*_(80)_ = 4.52, *p <* 0.0001, Bonferroni corrected), it never dropped to chance level (Figure 2B). On average, we found above chance decoding in both conditions in the early delay (Load 1: *t*_(80)_ = 11.71, *p <* 0.0001, Load 2: *t*_(80)_ = 9.00, *p <* 0.0001, uncorrected) and late delay (Load 1: *t*_(80)_ = 7.50, *p <* 0.0001, Load 2: *t*_(80)_ = 5.61, *p <* 0.0001, uncorrected).

Frontoparietal regions were not affected by load late in the delay. In IPS, our analysis revealed main effects of load (*F*_(1,80)_ = 7.07, *p* = 0.009) and time (*F*_(1,80)_ = 7.81, *p* = 0.006), as well as an interaction between load and time (*F*_(1,80)_ = 5.39, *p* = 0.023) on decoding accuracy. Here, decoding accuracy was reduced in visual Load 2 compared to Load 1 early, (*t*_(80)_ = *−*3.50, *p* = 0.002, Bonferroni corrected), but not late in the delay (*p* = 1.0, Bonferroni corrected). There was a significant difference in the change in decoding accuracy across time between load conditions. While the decoding accuracy for representations in the visual Load 2 condition showed no difference between the early and late delay period in IPS (*p* = 1), the decoding accuracy for the single visual item dropped from early to late delay (*t*_(80)_ = *−*3.29, *p* = 0.004, Bonferroni corrected). There was above chance decoding in both conditions in the early delay (Load 1: *t*_(80)_ = 7.03, *p <* 0.0001, Load 2: *t*_(80)_ = 4.29, *p <* 0.0001, uncorrected) and late delay (Load 1: *t*_(80)_ = 4.32, *p <* 0.0001, Load 2: *t*_(80)_ = 3.62, *p* = 0.0005, uncorrected).

In sPCS, while there was above chance decoding in both delay periods and conditions, there was no significant main effects or interaction effects of load conditions and delay period (all *p >* 0.7). There was no difference in decoding accuracy between load conditions in either early (*p >* 0.8, Bonferroni corrected) or late delay (*p >* 0.8, Bonferroni corrected). There was no difference between early and late delay for either Load 1 (*p >* 0.9) or Load 2 (*p >* 0.6, Bonferroni corrected).

### 3.2 Decoding accuracy correlates with behavioural recall precision

Our results so far indicate that anterior as well as posterior regions store working memory representations. To quantify the contributions of the three regions to behavioural performance in each load condition, we correlated late delay decoding accuracy and behavioural precision across all participants. Overall decoding accuracy correlated positively with behavioural precision in V1-V3 (*r* = 0.31, *p* = 0.005),IPS (*r* = 0.22, *p* = 0.046) and sPCS (*r* = 0.26, *p* = 0.018). Decodable information of orientations in visual cortex during the late delay correlated with behavioral performance in both visual load conditions (*r*_*Load*1_ = 0.24, *p* = 0.028, *r*_*Load*2_ = 0.25, *p* = 0.023). However, behavioural precision correlated significantly with decoding accuracy in IPS only in the Load 2 condition (*r* = 0.27, *p* = 0.013, Figure 2D), but not in the Load 1 condition (*r* = 0.11, *p* = 0.32). In sPCS, decoding accuracy correlated overall with behavioural precision, but not significantly for either Load 1 (*r* = 0.11, *p* = 0.35) or Load 2 (*r* = 0.22, *p* = 0.054).

Further analysis comparing the overall correlation coefficients for each ROI and behavioural precision show that they are not significantly different from each other (all comparisons *p >* 0.4). However, it is possible that the decoding accuracy in the other regions influences these correlations. When controlling for the effects of IPS and sPCS, a partial correlation analysis indicated a unique contribution of decoding accuracy in the visual cortex to behavioral precision (*r* = 0.24, *p* = 0.029).

### 3.3 Simulations indicate different information content in visual versus anterior regions

Our results indicate that in visual cortex, but less so in frontoparietal areas, decoding accuracy for orientations is reduced when multiple items are maintained in working memory. But is our pSVR analysis able to differentiate overlapping representations and quantify the strength of the underlying representations? We simulated multivariate patterns with our experimental parameters and compared a Load 1 condition to different variants of a Load 2 condition (see Methods and Figure 3A and 3B).

To determine how our decoding scheme impacts the results under varying signal strength, we simulated a Load 1 condition, where only one item is held in mind and two Load 2 conditions, one where the items were decoded separately and one where the items were decoded jointly, as in our analysis. In Figure 3C we show that with increasing signal strength, FCA increases. However, in both decoding schemes for the Load 2 conditions, the maximal decoding accuracy is around 25% above chance, while items in Load 1 can be hypothetically decoded with almost 50% above chance (i.e., 100% accuracy). This shows that if both items are maintained in overlapping patterns, even in data where the signal is weighted 10 times more than the noise, the data itself limits the hypothetical maximum decoding accuracy: the decoder has to select one of the two possible labels associated with this multivariate pattern. However, when the signal decreases, under conditions similar to our experimental data, we show that the load conditions do not carry over this drastic difference. Instead, the decoding schemes achieve similar decoding accuracies for both load conditions, where the joint decoding increases sensitivity slightly compared to the separate decoding.

In this scenario, we assume that in the Load 2 condition both items are maintained in the multivariate signal and thus a capacity of two (Load 2-Capacity 2). However, it might be possibly that visual cortex only maintains one of the two items (Sutterer et al., 2019), perhaps through a similar mechanism previously implicated in the prioritization of working memory items. When two items were maintained, only the one relevant for the upcoming task was reliably decodeable from visual cortex. Both items, the one immediately relevant and the item relevant for a future task were maintained in frontoparietal areas (Christophel et al., 2018). To test this possibility, we simulated a Load 2 condition where on each trial one of the two orientations was randomly selected to be maintained, thus simulating a capacity of one (Load 1-capacity 1; see also Sutterer et al. (2019)). While we varied the signal strength overall, we kept the strength of concurrent items the same in all simulations.

Our results for the comparison between Load 1 and Load 2-capacity 2, where the two patterns overlap, show that there is only a minor drop in FCA. This indicates that we can indeed decode information about orientations from overlapping patterns, if this drop is accounted for. On the other hand, FCA dropped substantially when we simulated a capacity of 1 (Load 2-capacity 1, Figure 3C), indicating that a loss of information in the Load 2 condition can be identified with our methods, under signal conditions similar to our experimental data.

How do these simulations compare to our empirical findings? We matched the simulated signal strength for each empirical Load 1 condition in each ROI and compared the ratio of FCA drop from Load 1 to Load 2 of our simulated results to our empirical results (Figure 3D and Methods). In the parietal and frontal cortex, our empirical results show high similarity to simulated results of a capacity of 2, where both items are concurrently maintained in working memory. This suggests that there is little cost to storing additional visual items in these regions. In visual cortex, empirical FCA dropped below the level expected when two orientations would be maintained (Load 2-capacity 2), but was higher than the FCA expected when only one of the items would be maintained (Load 2-capacity 1), suggesting that more than one visual item can be maintained in V1-V3, but at a cost for the individual representations.

### 3.4 Shared representations in visual cortex, distinct mnemonic code in sPCS

Considering that we observed changes in decoding accuracy across delay phases, we tested whether the neural code changes across time. First, we probed whether the multi-voxel pattern at one time-point generalizes to other time-points, within each load condition. Importantly, cross-decoding scales with the overall decoding accuracy, which limits the interpretability of these results.

When a single orientation (Load 1) is maintained in memory, the neural codes seemed to generalize between early and late delay time-points in all regions but sPCS(Figure 4A). In the Load 2 condition, temporal generalization is reduced, in accordance with the lower decoding accuracy early in V1-V3 and IPS (Figure 4B). Temporal generalization in sPCS was limited to neighboring time-points later in the delay, potentially due to limitations in power.

In visual cortex, we find evidence for dynamic clusters, where off-diagonal accuracy is lower than both diagonal time-points, for both load conditions. The neural code in visual cortex seemed to be changing dynamically from perception to the maintenance period. The IPS showed fewer dynamic clusters, while no dynamic clusters were found in sPCS.

Next, we tested whether the neural code for orientations is shared between load conditions. We trained the pSVR on each time-point in one load condition and tested on each time-point in the other load condition. In visual and parietal cortex, we found significant cross-condition decoding (diagonal in Figure 4C and D), suggesting that these representations generalized well across conditions. In sPCS, cross-condition decoding appears to be lower than within-condition decoding.

Overall, these results suggest that cortical working memory representations generalize across time, but not without loss of representational precision. It is further possible that the low cross-condition decoding in sPCS indicates different neural codes used for mnemonic representations in the two load conditions in this regions.

## 4 Discussion

Visual working memory representations are distributed across the cortex (Christophel et al., 2017). We investigated the effect of working memory load on these distributed representations by keeping the overall load constant at two items, but varying the sensory modality of each item. Our results suggest that the precision of visual working memory recall is limited by the resources available in visual cortex for the maintenance of multiple similar items. In contrast, increasing visual working memory load did not reduce item-specific information in IPS and sPCS during working memory maintenance. The item-specific decoding accuracy of the late delay period in all three regions predicted behavioural recall precision. Our cross-decoding and temporal generalization analyses hint at dynamic representational changes from perception to working memory in visual cortex and possibly load-dependent mnemonic representations in frontal regions.

Increasing visual working memory load reduced the ability to decode neural representations, as observed in previous studies (Emrich et al., 2013; Sprague et al., 2014; Sutterer et al., 2019). Here we show that for orientation stimuli the decoding accuracy decreases early in IPS and during the whole delay period in V1-V3, and that information in all three regions (V1-V3, IPS and sPCS) correlates with behavioural precision across participants. Previous work suggests that information for motion direction declines with load in sensory areas and correlates with behavioural performance (Emrich et al., 2013). When increasing visual working memory load for spatial locations, occipital and posterior parietal areas showed a decrease in strength of neural representations, which was also reflected in the decrease in behavioural performance (Sprague et al., 2014). These studies, as well as our task, encouraged the precise recall of low-level visual features and thus precise neural encoding in visual areas. However, in our data, the representations in visual cortex seem to be affected by interference between items, resulting in reduced decoding and reduced behavioural precision. Simulations of multivariate pattern for one and two orientations (see also Sutterer et al. (2019)), suggest that while both orientations (Load 2) are maintained in working memory signals in visual cortex, it comes with a loss of representational strength.

On the other hand, frontoparietal memory representations were not disrupted by increased visual working memory load. This is in line with prior studies suggesting that parietal representations might support working memory maintenance in the face of increasing cognitive demand. For instance, while representations in the visual cortex can be disrupted by taxing perceptual distractors, those in IPS may be less affected (Bettencourt & Xu, 2016; Rademaker et al., 2019). When participants maintained two items but prioritized only one at a time, both items were decodable from IPS and sPCS, whereas only the prioritized item was reliably decodable from the visual cortex (Christophel et al., 2018). However, parietal representations were disrupted if the distractor was behaviourally relevant (Hallenbeck et al., 2021) or when multiple spatial locations were maintained (Sprague et al., 2014), as purely spatial representations likely rely more on parietal regions (Jerde et al., 2012). Perhaps these regions can maintain multiple items without the same loss in precision seen in visual areas, as their representations are more abstract and less precise (Chunharas et al., 2023). This would enable the maintenance of multiple items in distinct neural patterns, rather than depending on the same neural populations to represent detailed orientation information as observed in visual cortex, and thus reduce interference in frontoparietal areas. Our task required the maintenance of two items in each trial and frontoparietal regions have been implicated in maintaining modality independent mnemonic representations (Spitzer & Blankenburg, 2012). Thus, another possibility is that the resource is similarly distributed across the visual and auditory item (visual Load 1) as for both visual items (visual Load 2) condition, although there is little overlap between parietal regions for pitch information and orientation information (Czoschke et al., 2021).

Our results show fewer dynamic clusters and lower cross-decoding between load conditions in these frontoparietal regions. Previous studies have shown suggested dynamic population coding in primate prefrontal areas could maintain stable working memory representations (Murray et al., 2017; Spaak et al., 2017). In humans, visual areas appear to show stronger dynamic change in representations compared to frontoparietal areas when a single spatial location is maintained in working memory (Li & Curtis, 2023). However, the temporal generalization and cross-decoding results are confounded by the strength of decoding accuracy in each region. While decoding accuracy is not comparable across ROIs, as two-way decoding of working memory contents showed a lower base-rate in frontal areas compared to visual cortex (Bhandari et al., 2018), it is possible that decoders can pick up on the cortical retinotopy in visual cortex for orientations from multivoxel patterns (Freeman et al., 2011), but possibly not the abstract features stored by frontal neurons (Sreenivasan et al., 2014). If orientations are stored in a more abstract format in frontoparietal areas and even visual areas during the working memory delay (Chunharas et al., 2023; Duan & Curtis, 2024; Kwak & Curtis, 2022; Yan et al., 2023), our decoding method, which assumes a continuous feature space, would thus underestimate the amount of information decodable from the neural patterns. Further studies should seek to disentangle the effects of decoding information with different neural codes.

## Data and Code Availability

Data and code are available to researchers upon request. The data cannot be shared publicly, because participants did not provide informed consent for public sharing.

## Author Contributions

VC and TC designed the study. VC collected the data. VC and TC performed data analysis. SW contributed software and materials. VC and TC wrote the manuscript.

## Acknowledgements

We thank Andreea Gui, Damla Cifci, Joana Seabra and Zhiqi Kang for their support with the fMRI data collection and the participants for their participation in this study. We thank Andreea-Maria Gui and Myriam Hamon for their work on the simulation toolbox and Karolis Degutis for sharing his code for the cluster analysis. We thank Alessandra Souza, Christoph Bledowski, Philipp Deutsch and Rosanne Rademaker for their helpful input in discussions about the experimental design and data analysis. We thank Renata Cruz for comments on this manuscript.

